# Remdesivir does not affect mitochondrial DNA copy number or deletion mutation frequency in aged male rats

**DOI:** 10.1101/2022.07.10.499455

**Authors:** Allen Herbst, Solbie Choi, Austin N. Hoang, Chiye Kim, Diana Martinez Moreno, Debbie McKenzie, Judd M. Aiken, Jonathan Wanagat

**Affiliations:** Department of Agricultural, Food and Nutritional Sciences, University of Alberta, Edmonton, Canada; Department of Medicine, Division of Geriatrics, UCLA, Los Angeles, USA; Department of Biological Sciences, University of Alberta, Edmonton, Canada; Veterans Administration Greater Los Angeles Healthcare System, Los Angeles, USA

**Author notes:** Correspondence: Jonathan Wanagat, MD, PhD.

**Keywords:** remdesivir, mitochondrial DNA, genetics, aging, heart, kidney, skeletal muscle

## Abstract

Remdesivir is a leading therapy in patients with moderate to severe coronavirus 2 (SARS-CoV-2) infection; the majority of whom are older individuals. Remdesivir is a nucleoside analog that incorporates into nascent viral RNA, inhibiting RNA-directed RNA polymerases, including that of SARS-CoV-2. Less is known about remdesivir’s effects on mitochondria, particularly in older adults where mitochondria are known to be dysfunctional. Furthermore, its effect on age-induced mitochondrial mutations and copy number has not been previously studied. We hypothesized that remdesivir adversely affects mtDNA copy number and deletion mutation frequency in aged rodents. To test this hypothesis, 30-month-old male F333BNF1 rats were treated with remdesivir for three months. To determine if remdesivir adversely affects mtDNA, we measured copy number and mtDNA deletion frequency in rat hearts, kidneys, and skeletal muscles using digital PCR. We found no effects from three months of remdesivir treatment on mtDNA copy number or deletion mutation frequency in 33-month-old rats. For the 33-month-old control rats, the average mtDNA copy number per nucleus was 2567, 1100, and 1869 for heart, kidney, and quadriceps, respectively. MtDNA deletion mutation frequency was 2.6×10^−4^, 1.6×10^−4^ and 4.7×10^−3^for heart, kidney, and quadriceps, respectively. These data support the notion that remdesivir does not compromise mtDNA quality or quantity at old age in mammals. Future work should focus on examining additional tissues such as brain and liver, and extend testing to human clinical samples.

## Introduction

Coronavirus disease 2019 (COVID-19), caused by the severe acute respiratory syndrome coronavirus 2 (SARS-CoV-2) has been a pandemic for over two years [1, 2]. Older persons are at increased risk for hospitalization or death from COVID-19 [3, 4]. According to a modeling study, the rate of COVID-19 patients hospitalized increased with age: 1% for those 20 to 29 years old, 4% for those 50 to 59 years old, and 18% for those older than 80 years of age [5]. Moreover, the risk for death among individuals 80 years and older is 20-fold higher than among individuals 50 to 59 years old [6]. According to the Center for Disease Control, more than 75% of COVID-19 deaths have been in older patients, including more than a quarter in people aged 85 and older (https://covid.cdc.gov/covid-data-tracker/#datatracker-home).

Remdesivir (Veklury®) is indicated for the inpatient and outpatient treatment of COVID-19 and is standard of care for hospitalized patients with moderate to severe COVID-19. Approximately four million individuals have been hospitalized with COVID as of January 2022. Of these, up to ∼50% have received remdesivir [7]. Therefore, approximately two million US patients have received remdesivir thus far – a number that will increase given remdesivir’s recently expanded use as an outpatient therapy. Despite its approval by the Food and Drug Administration (FDA), however, limited clinical data exists pertaining to its safety [8]. Remdesivir has not been evaluated for geriatric use [9]. There is a knowledge gap regarding the safety of drugs such as remdesivir in geriatric populations as individuals >64 years of age are routinely underrepresented in clinical trials [10]. This gap requires research as older individuals are often the primary target population for drugs such as remdesivir.

Remdesivir is a prodrug that is converted *in vivo* into an adenosine nucleoside analogue and subsequently phosphorylated to its triphosphate activated form. The targeted activity of remdesivir is the inhibition of viral RNA-dependent RNA polymerase necessary for the replication of the viral genome [8, 11]. While remdesivir does not appear to have off-target effects on the mitochondrial RNA polymerase activity *in vitro* [11], as an ATP-analogue, the active metabolite of remdesivir can participate in numerous unexplored cellular and mitochondrial processes. For example, the active metabolite could affect ATP buffering by creatine kinase and myokinase interfering in muscle metabolism similarly to beta-guanidinoproprionic acid, an analog of phosphocreatine [12, 13]. Remdesivir could also inhibit the adenine nucleotide translocator (ANT), which exchanges ATP and ADP across the mitochondrial inner membrane. Genetic defects in ANT lead to mtDNA deletion mutations and mitochondrial myopathy [14]. MtRNA polymerase activity is critical in priming mtDNA synthesis and is thought to be a regulatory step [15]. Other nucleoside analogues such as azidothymidine (AZT) have serious mitochondrial side effects such as lactic acidosis and myopathy [16, 17].

To date, few studies have investigated the effects of remdesivir on mitochondria, despite nucleoside analogs typically being associated with mitochondrial toxicity [18]. Direct *in vitro* effects of remdesivir on isolated pig brain mitochondria showed little or no effect on mitochondrial respiration [19]. However, remdesivir induced persistent mitochondrial changes in an *in vitro* human cardiac stem cell model [20]. At high doses, remdesivir was toxic to mitochondria in two human intestinal and liver cell lines [21]. Remdesivir impedes mitochondrial DNA polymerase gamma activity but stimulates exonucleolytic activity [22]. Despite this, remdesivir did not alter mitochondrial DNA copy number in human liver cells [23] or skeletal muscle fibroblast cells, but increased mtDNA copy number in human neonatal dermal fibroblasts [22]. Higher doses of remdesivir in rhesus monkeys resulted in a significant loss of mtDNA [24] in an unspecified tissue. While these studies suggest that remdesivir generally does not impact the quality or quantity of mitochondria, the *in vivo* effects of remdesivir on mitochondrial DNA copy number and mutations in older animals is unknown. This is especially relevant as remdesivir is administered predominantly to older COVID-19 patients.

The expanded use of remdesivir has led to the identification of important side effects including cardiac and renal effects. In the cardiovascular system, remdesivir increased the risk of bradycardia (Reporting Odds Ratio 1.63) and hypotension [25]. In the renal system, remdesivir treatment can cause acute renal failure, with 65-74-year-old old adults being at the highest risk [26, 27]. The role for mitochondria in these side effects is unclear but is likely related age-induced changes in these tissues and the reliance of heart and kidney on oxidative metabolism. Musculoskeletal side effects have not been reported for remdesivir treatment of COVID-19 patients.

MtDNA copy number and deletion frequency predict age and some measures of physical function in older adults [28, 29] and may be metrics of biological age. We hypothesized that remdesivir would negatively affect mtDNA quality by decreasing copy number and increasing mitochondrial mutation frequency in aged animals. To test this hypothesis, we measured mtDNA copy number and deletion mutation frequency in heart, kidney, and skeletal muscle from 33-month-old male rats treated with remdesivir for 3 monthsin. Remdesivir had no effect on either mtDNA copy number or mutation frequency at old age in this model, indicating that remdesivir does not negatively impact *in vivo* mitochondrial DNA quality and quantity in aged animals.

## Materials and methods

### Animals, remdesivir treatment, and tissue preparation

This study was carried out in accordance with the recommendations in the NIH Guide for Care and Use of Laboratory Animals and the guidelines of the Canadian Council on Animal Care using protocols approved by the Institutional Animal Care and Use Committees at UCLA and the University of Alberta. Thirty-month-old male Fischer 344 x Brown Norway F1 hybrid rats were obtained from the NIA Aging Rodent Colony. Remdesivir (MW 602.6) was purchased from VulcanChem (Pasadena, CA) and its authenticity was confirmed by mass spectrometry (data not shown). Drug delivery was via the subcutaneous implantation of a ninety-day time-release pellet containing 200 mg of remdesivir (Innovative Research of America, Sarasota, FL). The dose delivered over the 90 days was approximately 4.25 mg/kg/day. Rats were housed on a 12-hour light/dark cycle and fed standard chow. Control rats were implanted with a placebo pellet prepared from the time-release matrix only. Animals were euthanized by carbon dioxide asphyxiation followed by exsanguination. Tissues were dissected from the rats, weighed, and flash frozen in liquid nitrogen, and stored at -80°C.

### DNA isolation

Rat heart, kidney, and quadriceps muscle were ground to a powder using a mortar and pestle under liquid nitrogen. Total DNA was extracted using proteinase K digestion with SDS and EDTA, phenol/chloroform extraction, and ethanol precipitation. Total DNA was resuspended in 10 mM Tris-EDTA buffer, pH 8. Total DNA quality and quantity was assessed using spectrophotometry at A230, A260, and A280 (ThermoScientific Nanodrop 2000 Spectrophotometer), fluorometry (ThermoFisher Qubit 2.0 Fluorometer) and agarose gel electrophoresis.

### MtDNA copy number and mtDNA deletion frequency by digital PCR

A 5-prime nuclease cleavage assay and droplet-based digital PCR (dPCR) were used to quantitate copy numbers for nuclear DNA (nDNA), total mtDNA, and mtDNA deletion mutations, with specific primer/probe sets for each as previously described [30]. MtDNA deletion frequency is the proportion of mutant molecules per wild-type mtDNA. Deletions per 100 nuclei is a value normalized to the single copy nuclear gene in the rat, Unc13 [31]. DPCR quantitation of all samples and all targets was performed on coded samples. Blinding was removed following data collection.

## Statistical analysis

All data are presented as means ± SEM. Data were tested for a normal distribution. Student’s t-test was used to compare differences between treatment and control groups. Chi-squared test was used to determine differences in survival frequency. One-way analysis of variance was used to test statistical differences between tissues. Prism (Version 7.05, GraphPad Software) was used for all statistical analyses.

## Results

### Husbandry and morphometric measures following remdesivir treatment

Ninety days of subcutaneous remdesivir treatment starting at 30 months in male F344BN F1 hybrid rats had no adverse effects on food consumption (data not shown). The mean lifespan of male F344BNF1 rats is 34 months [32] and rat survival was not affected by remdesivir treatment with three control and two remdesivir-treated rats reaching a point where rat body condition scoring indicated that death was imminent and animals were culled during the 90 days of the experiment (Chi squared = 0.582). Body, heart, kidney, and muscle weights also were not affected by the remdesivir treatment (**Fig. 1**).

**Fig 1.**
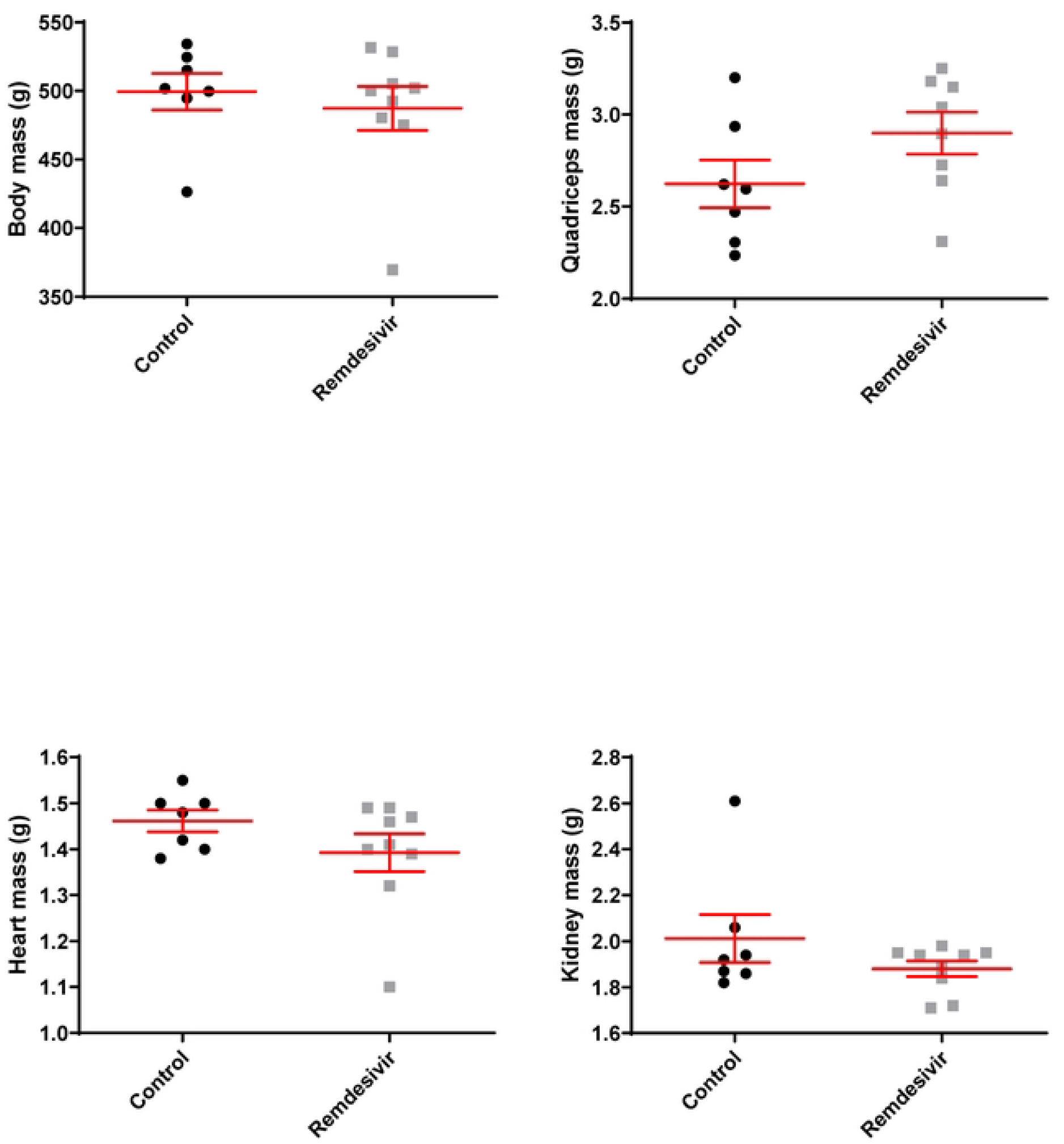
Effects of three months of remdesivir treatment on body, heart, kidney and quadriceps mass in hybrid rats. Rats were weighed at 33 months of age before they were sacrificed. Whisker plots denote mean and SEM. Black circles denote control rats, grey squares denote remdesivir-treated rats. N=7-9 per experimental group.

### Measures of mtDNA copy number and deletion mutation frequency following remdesivir treatment

Similar to the husbandry and morphometric outcomes, mtDNA copy number and deletion mutation frequency were not affected by 90 days of remdesivir treatment (**Fig. 2**). To assess the cellular burden of mtDNA deletion mutations, we calculated the number of deletion mutations per diploid nucleus and found no effects of remdesivir. MtDNA copy number and deletion mutation frequency differ greatly between heart, kidney, and skeletal muscle at 33 months. Heart mtDNA copy number was 1.70-fold higher than kidney and 1.37-fold higher than quadriceps, while the deletion mutation frequency in skeletal muscle was 18-fold higher than heart and 29-fold higher than kidney (**Fig. 3**).

**Fig 2.**
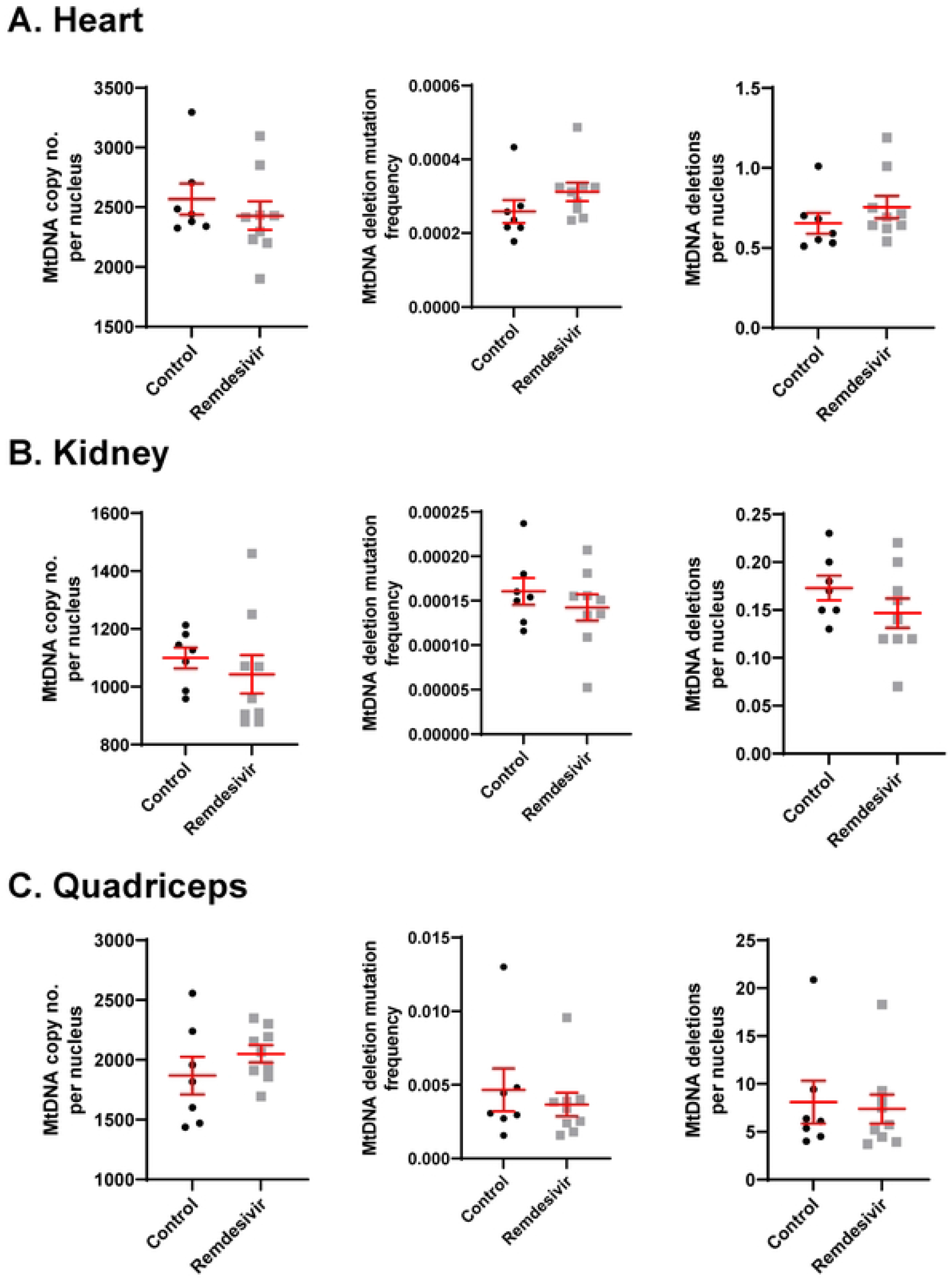
Effects of three months of remdesivir treatment on mtDNA copy number, deletion frequency, and deletions per nucleus in 33mo hybrid rats. A. heart, B. kidney, and C. quadriceps muscle. Whisker plots denote mean and SEM. Black circles denote control rats, grey squares denote remdesivir-treated rats. N=7-9 per experimental group.

**Fig 3.**
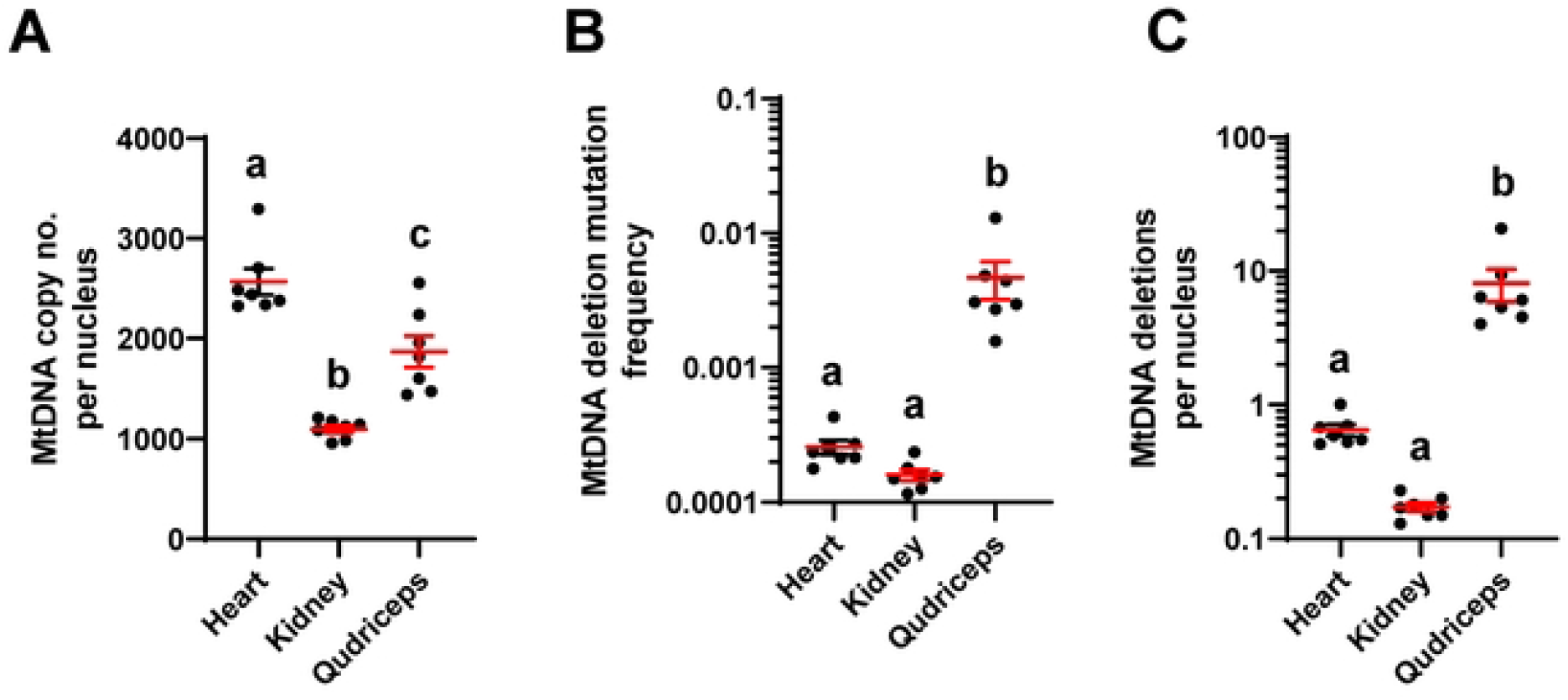
Tissue differences in A) mtDNA copy number, B) deletion mutation frequency, and C) deletions per nucleus in control 33mo F344BNF1 hybrid rats. Y-axes in B and C are log10 transformed. Whisker plots denote mean and SEM. Black circles denote control rats. N=7-9 per experimental group. Different lower case letters denote significance at p<0.05.

## Discussion

To our knowledge, this is the first study examining the impact of remdesivir on mtDNA metrics in aging rodents. As mitochondrial dysfunction is a conserved phenotype of aging with accessible and sensitive biomarkers, we investigated the effect of remdesivir on mitochondrial genetics, i.e., copy number and mtDNA deletion mutation frequency, in aged rats. We found that 90 days of remdesivir treatment did not alter mitochondrial copy number or deletion mutation frequency in 33-month-old male F333BNF1 rats. Furthermore, in the aged rats, we did not observe any effect on body weight, food consumption, or organ weights. Our data adds to the limited information available on remdesivir’s effect on mitochondria, especially in older individuals. Also, unlike other studies that focus on a single tissue, our study examined remdesivir’s impact on mitochondrial quality in multiple organ/tissue systems, specifically, heart, kidney, and skeletal muscle. Our findings indicate that the use of remdesivir to treat COVID in aging patients is not detrimental to their mitochondrial indices.

Remdesivir’s *in vivo* effects on the mtDNA polymerase and its subsequent impact on mitochondrial genetics are just beginning to be understood. Our data do not support the hypothesis that remdesivir affects mtDNA *in vivo*, despite the varied *in vitro findings* and *in vivo* findings in young animals [22, 23]. Moreover, although IV remdesivir in rhesus macaques, at a therapeutic concentration of 1μM remdesivir, showed no significant reduction in mtDNA copy number, a higher therapeutic level (2 μM) resulted in a 26% decrease in mtDNA copy number [24]. In contrast, when Bjork and Wallace [23] assessed the dose-dependent effect of remdesivir on mitochondrial DNA replication by exposing human hepatoma HepG2/C3A cells to increasing concentrations of remdesivir (0.1 to 10 μM) for 24 h and 48 h, they did not observe changes in mtDNA copy number. Given the pharmacokinetics of remdesivir in rats [33], our daily dose of 4.25 mg/kg of remdesivir is comparable to the human dose of 1.25mg/kg.

The effects of remdesivir on the accumulation of mtDNA deletion mutations has received very little attention. In human neonatal dermal fibroblasts, remdesivir treatment did not induce the 4977 or “common” mtDNA deletion mutation [22]. As a lower mtDNA copy number predicts a higher mtDNA deletion mutation frequency in human skeletal muscle [29], we hypothesized that reductions in mtDNA copy number induced by remdesivir would result in increased mtDNA deletion mutation frequency. Our data indicate that remdesivir does not contribute to age-induced mitochondrial genetic structural rearrangements.

Little is known about the effect of remdesivir on mitochondrial transcription. Bjork and Wallace [23] studied the dose-dependent effect of remdesivir on mitochondrial gene expression *in vitro* and found no evidence that remdesivir interferes with gene transcription. In a separate study, when HepG2 (liver) and HT-29 (intestinal) cells, treated with remdesivir for 8 and 24 h (in contrast to 24 and 48 h for Bjork et al), were analyzed by RNAseq, there was a decrease in the expression of genes involved in mitochondrial respiration and an observation that remdesivir decreases cellular ATP (Akinci, Cha et al. 2020). They hypothesized that this reduction in ATP was, at least in part, due to a reduction in mitochondrial gene expression. The discrepancy in these two studies should be tested, using aged rodents, in transcriptomic studies such as RNAseq. Aged rodent studies would have the added advantage of utilizing *in vivo* samples, compared to the aforementioned *in vitro* studies, providing a clearer and more accurate picture. Additionally, an aged rodent model would be conducive to testing remdesivir for side effects that might be expected in older adults.

Unlike human COVID-19 patients who are administered remdesivir via intravenous infusion, the aged rats in this study were given remdesivir in the form of a subcutaneously implanted time-release tablet. The remdesivir treatment in our study was considerably longer – 90 days – as opposed to a 5-10-day course in humans. We would, therefore, predict more pronounced side effects in the aged rats treated with remdesivir, compared to those seen in humans. This is not what we observe. Our results demonstrate that remdesivir is not detrimental to mitochondrial quality and quantity in an appropriately aged animal model. We examined rat heart, kidney, and skeletal muscle, but remdesivir has also been reported to have hepatic side effects [34]; we did not examine liver tissue for mtDNA effects.

In summary, our data demonstrate that remdesivir does not compromise mitochondrial copy number or mtDNA deletion mutation frequency in heart, kidney, and skeletal muscle in aged rodents. In addition to testing its impact on other relevant tissues like brain and liver, future work should focus on examining the effect of remdesivir on *in vivo* mitochondrial transcription. Finally, because rodent studies are unable to completely predict human outcomes, similar studies should be extended to clinical samples of human COVID-19 patients who received remdesivir treatment.

## Acknowledgements

Writing assistance was provided by Pranali Pathare Mangat, PhD of 3P Scientific Communications.

This material is the result of work supported with resources and the use of facilities at the Veterans Administration Greater Los Angeles Healthcare System.

